# Evolution of antiviral resistance captures a transient interdomain functional interaction between chikungunya virus envelope glycoproteins

**DOI:** 10.1101/2024.11.11.623010

**Authors:** Leandro Battini, Sara A Thannickal, Malena Tejerina Cibello, Mariela Bollini, Kenneth A Stapleford, Diego E Álvarez

## Abstract

Envelope proteins drive virus and host-cell membrane fusion to achieve virus entry. Fusogenic proteins are classified into structural classes that function with remarkable mechanistic similarities. Fusion proceeds through coordinated movements of protein domains in a sequence of orchestrated steps. Structures for the initial and final conformations are available for several fusogens, but folding intermediates have largely remained unresolved and interdependency between regions that drive conformational rearrangements is not well understood. Chikungunya virus (CHIKV) particles display heterodimers of envelope proteins E1 and E2 associated as trimeric spikes that respond to acidic pH to trigger fusion. We have followed experimental evolution of CHIKV under the selective pressure of a novel small-molecule entry inhibitor. Mutations arising from selection mapped to two residues located in distal domains of E2 and E1 heterodimer and spikes. Here, we pinpointed the antiviral mode of action to inhibition of fusion. Phenotypic characterization of recombinant viruses indicated that the selected mutations confer a fitness advantage under antiviral pressure, and that the double-mutant virus overcame antiviral inhibition of fusion while single-mutants were sensitive. Further supporting a functional connection between residues, the double-mutant virus displayed a higher pH-threshold for fusion than single-mutant viruses. Finally, mutations implied distinct outcomes of replication and spreading in mice, and infection rates in mosquitoes underscoring the fine-tuning of envelope protein function as a determinant for establishment of infection. Together with molecular dynamics simulations that indicate a link between these two residues in the modulation of the heterodimer conformational rearrangement, our approach captured an otherwise unresolved interaction.

**Importance:** CHIKV is a reemergent pathogen that has caused large outbreaks in the twenty years. There are no available antiviral therapies and a vaccine has only recently been approved. Here, we describe the mode of action of a novel inhibitor designed against CHIKV envelope proteins heterodimer that blocks entry at the stage of fusion between virus and host membranes. Fusion is common to the entry of enveloped viruses. Virus envelope proteins drive fusion undergoing a series of transitions from an initial metastable conformational state to a more stable post-fusion state. Intermediate conformations are transient and have mostly remained inaccessible to structure determination. In this study, directed evolution of resistance to antiviral inhibition of fusion uncovered a functional interaction between two residues residing in domains that are apart in both the pre-fusion and post-fusion states. Thus, our approach allowed gaining insight into the molecular detail of the inner working of virus fusion machinery.

## Introduction

Chikungunya virus (CHIKV) is transmitted to humans by mosquitoes. Infection commonly results in acute fever and joint pain that can progress into chronic polyarthritis (1,2). Since 2004, CHIKV has spread throughout the tropical and subtropical regions around the world causing outbreaks associated with high socioeconomic cost (3,4). Genomic adaptation of the virus to new mosquito vectors has been pointed out as one of the major causes for the geographical spread of the virus (1,5,6). Therefore, the study of viral determinants of CHIKV fitness in its natural hosts may provide valuable information regarding the continued adaptation of the virus.

CHIKV is an alphavirus of the *Togaviridae* family (7). It has a 12 kb positive stranded RNA genome that encodes for 10 proteins in two open reading frames (ORFs). The second ORF in the 3’ of the genome encodes for the structural proteins of the virus: the capsid (C) and the envelope glycoproteins (E3-E2-E1). E2 and E1 are transmembrane proteins that together with E3 form a heterotrimer in the surface of the viral particle, and mediate the interaction with cell receptors and fusion between viral and endosomal membranes during entry, respectively (8). E2 belongs to the immunoglobulin superfamily of proteins and folds into three globular domains (A, B and C) connected by two antiparallel β-strands, referred to as β-ribbon (9). E1 is a class II viral fusion protein and folds into a β-sheet rich structure with three β-barrel domains (I, II and III) bearing the fusion loop in the tip of domain II (9,10). In the viral particle, three E3-E2-E1 trimers fold into a viral spike, with E1 located next to the membrane and E2 positioned over E1 protecting the fusion loop (11).

CHIKV enters the cell by receptor-dependent endocytosis. In the final stage of the entry process, the decrease in endocytic pH induces membrane fusion triggering a major conformational rearrangement of the envelope proteins that involves dissociation of E1 and E2, and reassociation of E1 into homotrimers (9,12–15). The joint action of different E1 homotrimers is necessary to provide the driving force required to curve the opposing membranes (10,16). For this reason, the conformational rearrangement in different spikes must be coordinated and the timing of each step must be tightly regulated.

We have previously identified compound 11 as a small molecule inhibitor of CHIKV infection following a virtual screening against the envelope proteins of the virus (17). We showed that compound 11 inhibited the internalization of the viral particle and selected a resistant variant to the antiviral activity of the compound harboring mutations in E1 (Y24H) and E2 (P173S). Interestingly, recombinant viruses carrying both E1-Y24H and E2-P173S showed the resistant phenotype but neither of the single mutants did, suggesting a functional interaction between these two residues located in distant regions of CHIKV envelope glycoproteins (Figure 1).

**Figure 1:**
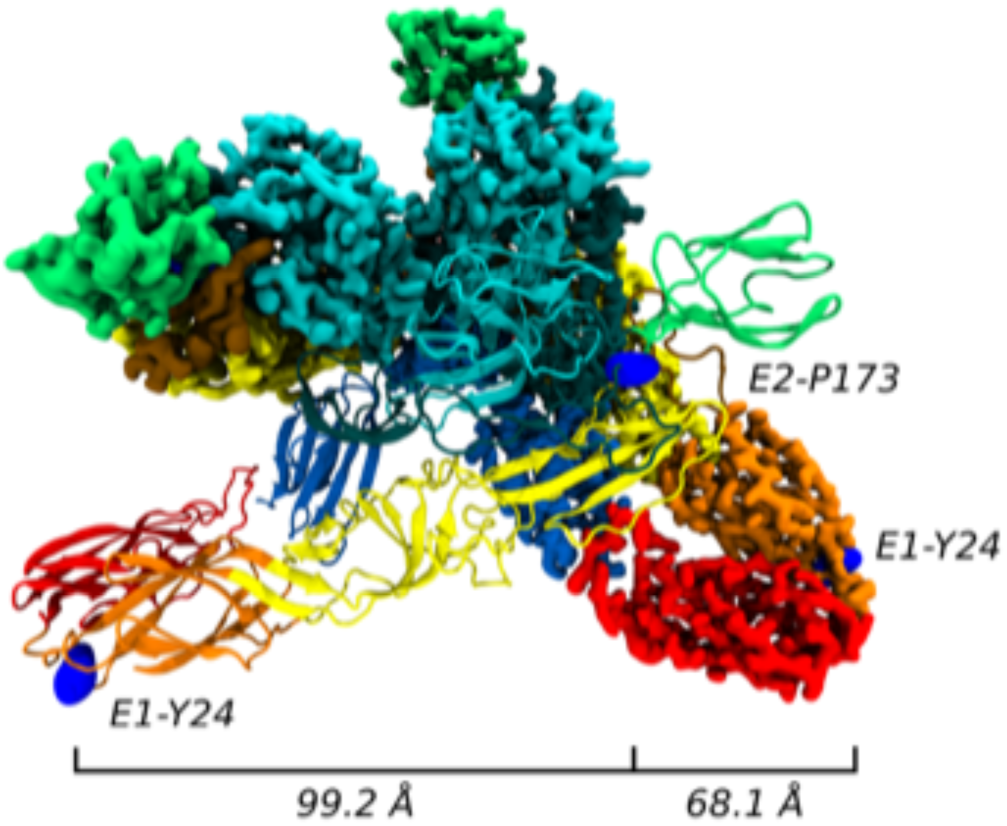
Mapping of E2-P173 and E1-Y24 on the trimeric spike. Representation of the spike formed by three E2-E1 heterodimers (PDB: 6NK7). One heterodimer is represented as a cartoon and the remaining two as surface. Domain E2-A is colored in cyan, E2-B in green, E2-C in blue, E1-I in orange, E1-II in yellow and E1-III in red. The β-ribbon is colored in dark cyan and the fusion loop in ochre. E2-P173 and E1-Y24 are represented as blue surfaces.

In this study, we show that compound 11 inhibits the pH-dependent membrane fusion process. The characterization of E1-Y24H and E2-P173S single and double mutant viruses showed that the double mutant displays an altered pH dependance for fusion that differs from both WT and single E1-Y24H and E2-P173S mutant viruses. Furthermore, the double mutant virus showed enhanced replication in mice but impaired ability to infect mosquitoes compared to WT virus. Based on molecular dynamics simulations of the pre-fusion conformation of the envelope proteins, we propose that residues E1-Y24 and E2-P173 are associated with kinetic barriers that act as checkpoints of the protein conformational change during the fusion process. Altogether, the results presented in this work support a functional interaction between residues E1-Y24 and E2-P173 that function in a concerted manner to regulate the fusion process.

## Results

### Compound 11 inhibits membrane fusion

We have identified a small molecule inhibitor of CHIKV infection. Time of drug addition assays showed that compound 11 inhibited the release of viral progeny when added at the time of infection and that the effect was gradually lost when addition was delayed indicating that treatment blocked virus entry but not the release of infectious particles. In line with this observation, we found that the compound had no effect on virus attachment and specifically inhibited internalization, reducing relative infection by 3-fold in Vero cells treated with 50 µM of compound 11 (17). To further characterize its mode of action, we tested the inhibitory activity of the compound against lentiviruses pseudotyped with CHIKV or VSV envelope proteins using BHK cells as a target for transduction (Figure 2A). While the VSV pseudotype was resistant to antiviral activity of compound 11, it inhibited CHIKV pseudotyped lentivirus (EC_50_ = 7.6 ± 0.1 μM, Figure 2A inset). The EC_50_ is similar to the value measured for compound 11 in BHK cells against a reporter CHIKV expressing ZsGreen from a subgenomic promoter to assess infection levels (EC_50_ = 13.77 ± 1.87 μM and Figure 2B), demonstrating that the compound specifically inhibits CHIKV entry into the host cell. Moreover, both CHIKV and VSV are endocytosed through the clathrin-dependent pathway (18,19). As we have shown that compound 11 did not interfere with virus attachment (17), the fact that inhibition was specific against the CHIKV pseudotype rules out an indirect effect of the compound in endocytosis and suggests that compound 11 may inhibit the function of CHIKV envelope proteins in the membrane fusion process. Then, we performed Fusion From Without (FFWO) assays in which fusion of viruses attached on the cell surface is triggered directly at the plasma membrane by lowering the pH of the medium. As a result, the degree of infection is dependent on the fusogenic activity of the viral particles. We used a reporter virus expressing ZsGreen from a subgenomic promoter to assess infection levels (Figure 2B). First, by triggering fusion at different pHs (Figure 2C) we observed that the fusogenic activity of the envelope proteins followed a sigmoidal curve with a fusion threshold around pH 5.5, similar to the behavior described in literature (20,21). We then evaluated the effect of adding compound 11 at increasing concentrations in FFWO triggered at pH 5.4, 5.5 or 5.6. Figure 2D shows that compound 11 inhibited fusion in a concentration-dependent manner and had a stronger effect at higher pH.

**Figure 2:**
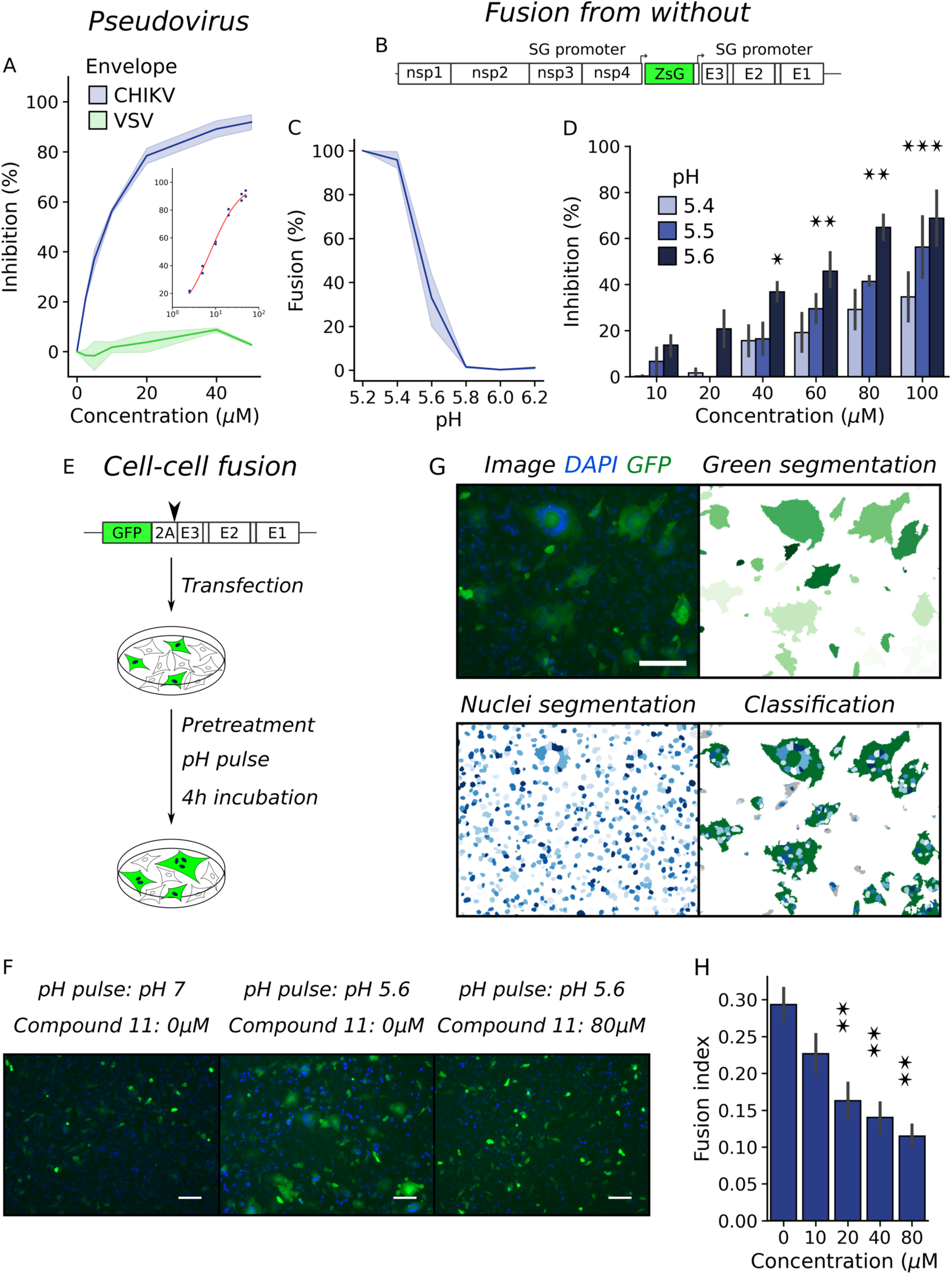
Compound 11 inhibits the fusion process during CHIKV entry. (A) Inhibition of pseudotyped lentivirus by compound 11 in BHK cells. Results represent the mean and standard error of two independent experiments. (B) Schematic representation of CHIKV ZsGreen genome. (C) FFWO assay with WT CHIKV ZsGreen virus in BHK cells triggering fusion with citric acid in PBS titrated at different pHs. Results represent the mean and standard error of the percentage of infected cells as determined by flow cytometry relative to the percentage at pH 5.2 of three independent experiments. (D) FFWO assay with CHIKV WT in the presence of compound 11. Results represent the mean and standard error of the inhibition of infected cells relative to the untreated control of four independent experiments. One way ANOVA with Tukey’s post hoc test. Differences against the untreated control for each pH, * p < 0.5. (E) Schematic representation of pCI-neo-CHIKV-GFP construct and workflow of cell-cell fusion assay. (F) Representative images obtained after different treatments in cell-cell fusion assay. The white scale bar represents 100 μm. (G) Image analysis of cell-cell fusion assay. From the original image, green cells and nuclei were segmented and green cells were classified in non-fused cells and syncytium. The fusion index was calculated as Fusion Index = 1 - number of green cells / number of nuclei. (E) Fusion index in the cell-cell fusion assay against increasing concentration of compound 11. Results represent the mean and standard error of three independent experiments. One way ANOVA with Tukey’s post hoc test , * p < 0.5, ** p < 0.01.

As a complementary approach, we set up a cell-cell fusion assay based on the ability of recombinant CHIKV envelope proteins exposed on the plasma membrane to induce fusion with target cells. In BHK cells transfected with a reporter construct encoding for GFP and CHIKV structural proteins separated by a self-cleaving peptide, green multi nuclei syncytia formed after treatment with low pH medium (Figure 2E-F). Fluorescence image analysis was followed to classify GFP expressing cells based on the number of encompassed nuclei, and the fusogenic activity was estimated using a fusion index (*Fusion Index = 1 - cells / nuclei*) that ranges from 1 to 0 with higher values indicating a higher fusogenic activity (Figure 2G). As in the FFWO assay, compound 11 inhibited the fusogenic activity of CHIKV envelope proteins in a concentration dependent manner when fusion was triggered at pH 5.6 (Figure 2H).

In conclusion, complementary approaches using pseudo typed viruses, fully infectious reporter CHIKV, and protein expression constructs to evaluate function of CHIKV envelope proteins demonstrate that compound 11 inhibited entry of CHIKV into the host cell and pinpoint the mechanism of action to the inhibition of the fusion process driven by CHIKV envelope proteins.

### E2-P173S E1-Y24H double mutant resistant phenotype is associated to overcoming of fusion inhibition

We next studied the resistant phenotype of E2-P173S and E1-Y24H double and single mutants. Recombinant viruses were previously constructed in the reporter CHIKV-ZsGreen background (17). First, growth curves in Vero cells showed that all viruses behaved in the same manner as WT (Figure 3A). To assess fitness in the presence of compound 11, we measured antiviral activity using reporter CHIKV-ZsGreen in a focus forming assay. WT and single mutants E2-P173S and E1-Y24H showed similar sensitivity to compound 11 displaying EC_50_ of 11.6 ± 3.4 μM, 15.7 ± 1.2 μM, and 11.2 ± 2.1 μM, respectively, while the double mutant displayed EC_50_ = 21.4 ± 1.3 μM, indicating that only the combined mutations confer a partially resistant phenotype to compound 11 antiviral activity. The phenotype was further confirmed when viral yields were measured following infection of cells treated with increasing compound concentrations (Figure 3B). In the absence of compound, the E2-P173 mutant yielded three-fold higher titers than WT, and E1-Y24H and double mutant viruses. As expected, increasing compound concentrations resulted in a gradual decrease in virus yields. While single mutants showed a trend similar to WT, E2-P173S E1-Y24H double mutant virus demonstrated a clear resistance to the antiviral activity of compound 11 displaying more than 100-fold higher viral yields at 25 μM of compound. The higher titers observed for the E2-P173S mutant compared to WT in the absence of compound were also observed across the concentrations tested, and together with the results in the focus forming assay indicate that the two viruses are similarly sensitive to the antiviral effect of compound 11.

**Figure 3:**
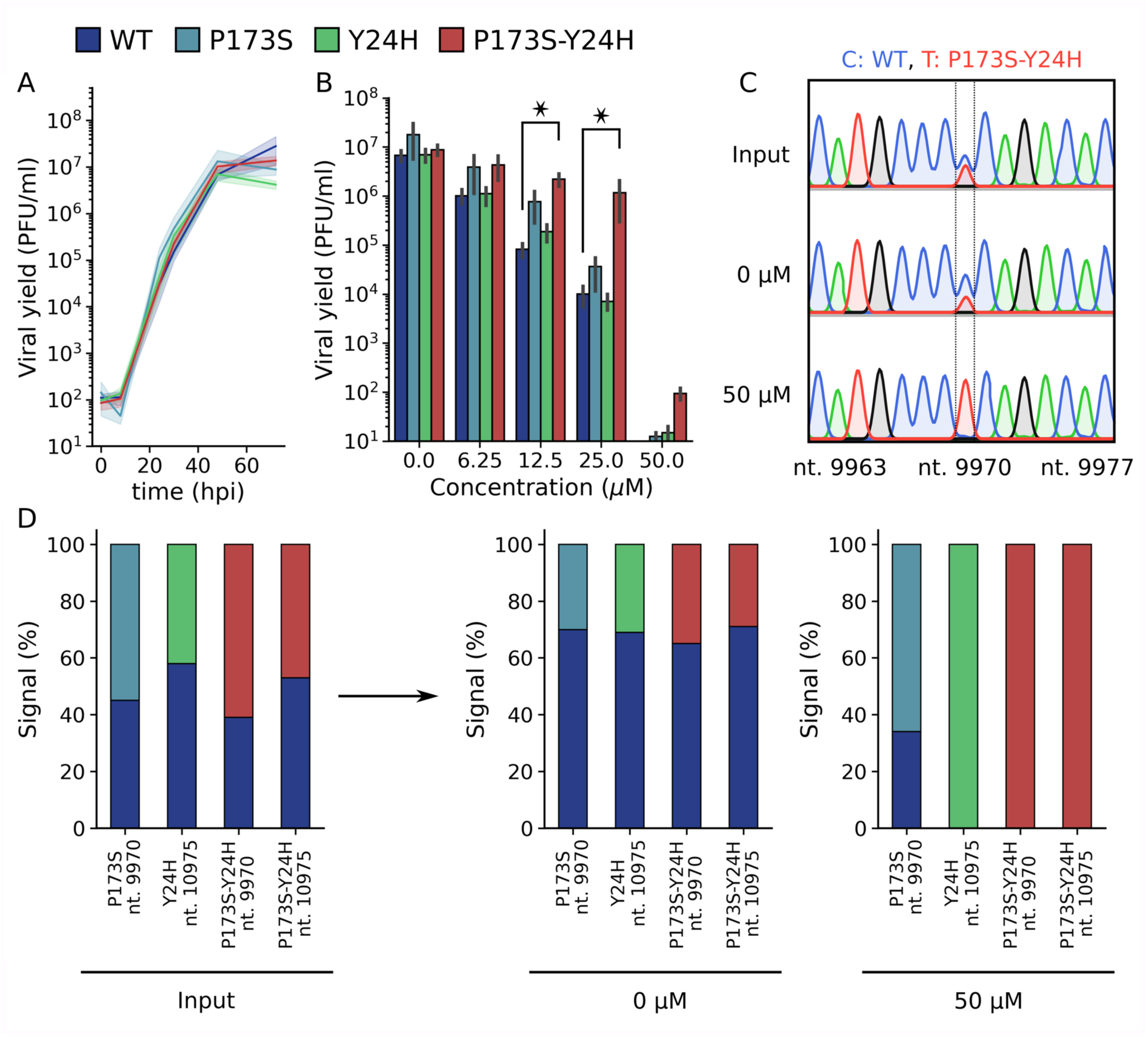
Single mutants E1-Y24H and E2-P173S, and double mutant viruses outcompete wild type virus growth in the presence of compound 11. (A) Growth curve of WT and mutant viruses on Vero cells in absence of compound. Results represent the mean and standard error of three independent experiments (B) Viral yield 48 after the infection with WT and mutant viruses and after the treatment with increasing concentrations of compound 11. Results represent the mean and standard error of three independent experiments. Kruskal-Wallis with Dunn’s post hoc test, * p < 0.05. (C) Representative chromatograms obtained in the competition assay. The nucleotide position is indicated in the x-axis label. Vero cells or cells treated at 50 μM of compound 11 were infected with WT and mutant viruses mixed at an initial 1:1 ratio, and the supernatants were collected at 72 hours after infection. Following RNA extraction and RT-PCR, the composition of the resulting virus population was assessed by Sanger sequencing. The peak height was used to determine the ratio between viruses. (D) Quantification of the competition assay. Signal percentages represent the ratio of the peak height for a given nucleotide divided by the sum of the heights of the nucleotides in the mixed signal. The nucleotide position is indicated in the x-axis label.

Next, the fitness of WT and mutant viruses was further evaluated in virus competition assays. Untreated cells or cells treated at 50 μM of compound 11 were infected with WT and mutant viruses mixed at an initial 1:1 ratio. Then, the composition of the resulting virus population was assessed by Sanger sequencing of the RT-PCR product amplified from cell culture supernatants by comparing the peak height of the WT and mutant alleles (Figure 3C). As expected, we observed a 1:1 signal ratio for input populations. In untreated cells, although mutant alleles were detected, in all cases the ratio favored the WT, suggesting that mutations impaired virus fitness. In turn, in treated cells, both E1-Y24H and the double mutant viruses completely displaced the WT virus, and the signal ratio for the E2-P173S and WT was inverted compared to competition in untreated cells, indicating that the mutant allele conferred an increment in fitness in the presence of compound 11 (Figure 3D). The association of individual mutations to resistance was not anticipated in focus forming and virus yield assays, but it was only evident in direct competition assays under stringent antiviral pressure. In turn, the assay also suggested that mutations have a cost in virus fitness when viruses competed without antiviral pressure suggesting that these conflicting forces would have acted together in the selection process that led to the emergence of the double mutant virus at high antiviral concentrations.

To gain insight into the mechanism by which the double mutant virus overcomes antiviral activity of compound 11, we evaluated the effect of compound 11 on infection with WT and single and double mutant pseudotyped lentiviruses (Figure 4A). At concentrations greater than 10 µM, only the double mutant virus displayed increased levels of infection compared to WT, indicating that the combined E2 P173S E1-Y24H mutations in the envelope proteins are sufficient to confer antiviral resistance.

**Figure 4:**
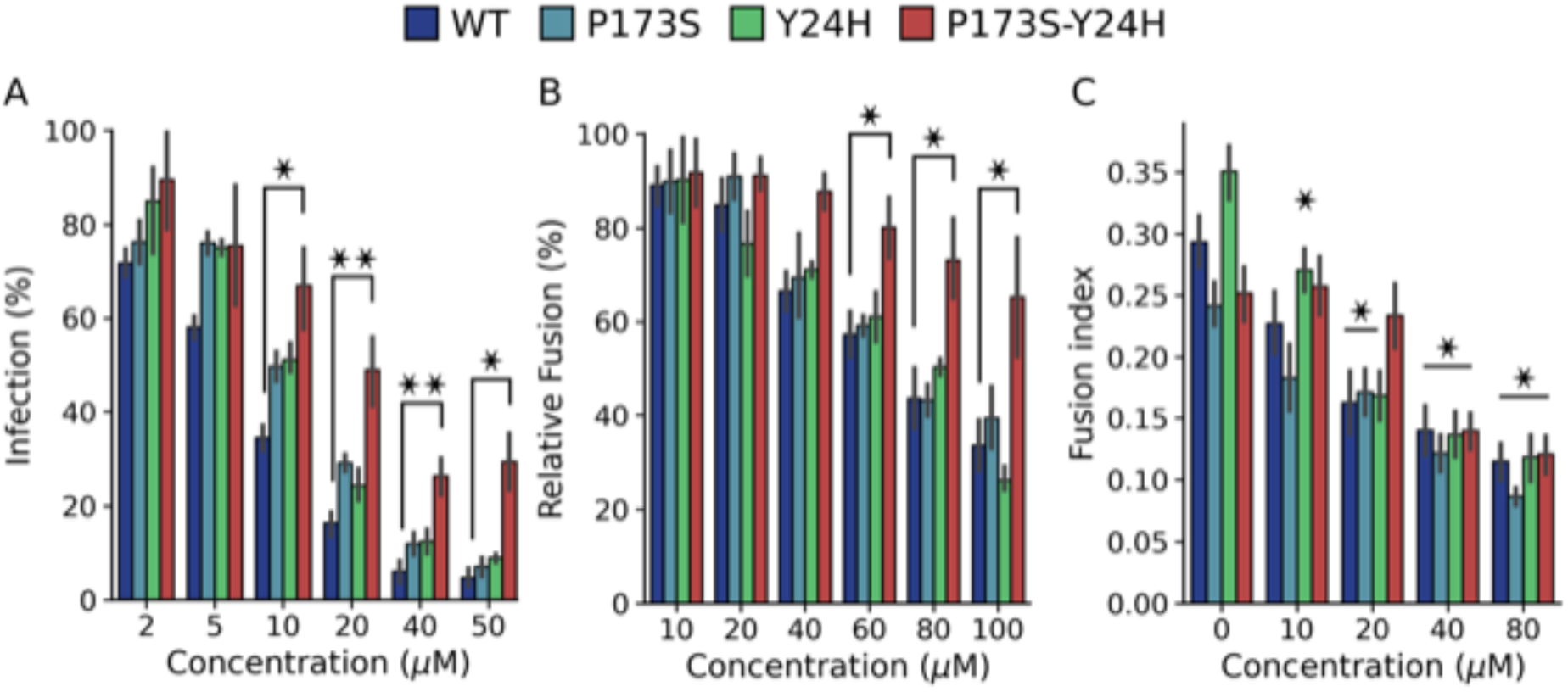
E1-Y24H E2-P173S double mutant escapes compound 11 fusion inhibition. (A) Inhibition of WT and mutant pseudotyped lentivirus with increasing concentrations of compound 11. Results represent the mean and standard error of the percentage of infected cells relative to the untreated control of three independent experiments. One way ANOVA with Tuekey’s post hoc test, * p < 0.05, ** p < 0.01. (B) Fusion inhibition of compound 11 in the FFWO assay (pH 5.6) with WT and mutant viruses. Results represent the mean and standard error of the percentage of infected cells relative to the untreated control of three independent experiments. One way ANOVA with Tukey’s post hoc test. * p < 0.05. (C) Fusion inhibition of compound 11 in the cell-cell fusion assay (pH 5.6). Results represent the mean and standard error of the Fusion Index of three independent experiments. One way ANOVA with Tukey’s post hoc. Differences between each concentration and the untreated control for each virus, * p < 0.05.

Next, we performed FFWO and cell-cell fusion assays with the WT and single and double mutant viruses to determine if the mutations allow the virus to overcome fusion inhibition by compound 11. In the FFWO assay, the single mutants behaved in the same manner as the WT virus showing a dose-dependent inhibition of infected cells (Figure 4B). In contrast, the double mutant infected a higher percentage of cells in the range of concentrations tested. In line with these results, in the cell-cell fusion assay, the fusogenic activity of the WT and both single mutants decreased as drug concentrations increased (Figure 4C). Contrarily, there were no differences in the fusogenic activity of the double mutant up to 20 μM of compound. Compared to the FFWO assay, resistance of the double mutant virus to the inhibitory effect in the cell-cell fusion assay was not observed at the highest concentrations tested likely due to methodological differences between approaches based on infection with replicative viruses and overexpression of envelope proteins, respectively.

Altogether, these results show that resistance of E2-P173S E1-Y24H double mutant to the antiviral activity of compound 11 is associated with a weaker inhibition of the fusogenic activity of CHIKV envelope proteins reinforcing the notion of a functional interaction between E2-P173 and E1-Y24.

### Mouse and mosquito infections with E2-P173S and E1-Y24H single and double mutants

To characterize the impact of mutations *in vivo*, we performed infections in mice with the WT and mutant viruses. Two days post-infection we quantified footpad viral titers that reflect virus replication at the site of infection, and viremia as a proxy of dissemination (Figure 5A and B). Footpad viral titers were higher for mutant viruses than for the WT with the double mutant displaying the highest viral yield. In this line, viremia was also higher for mutant viruses showing a positive correlation with the infection level in the primary infection site.

**Figure 5:**
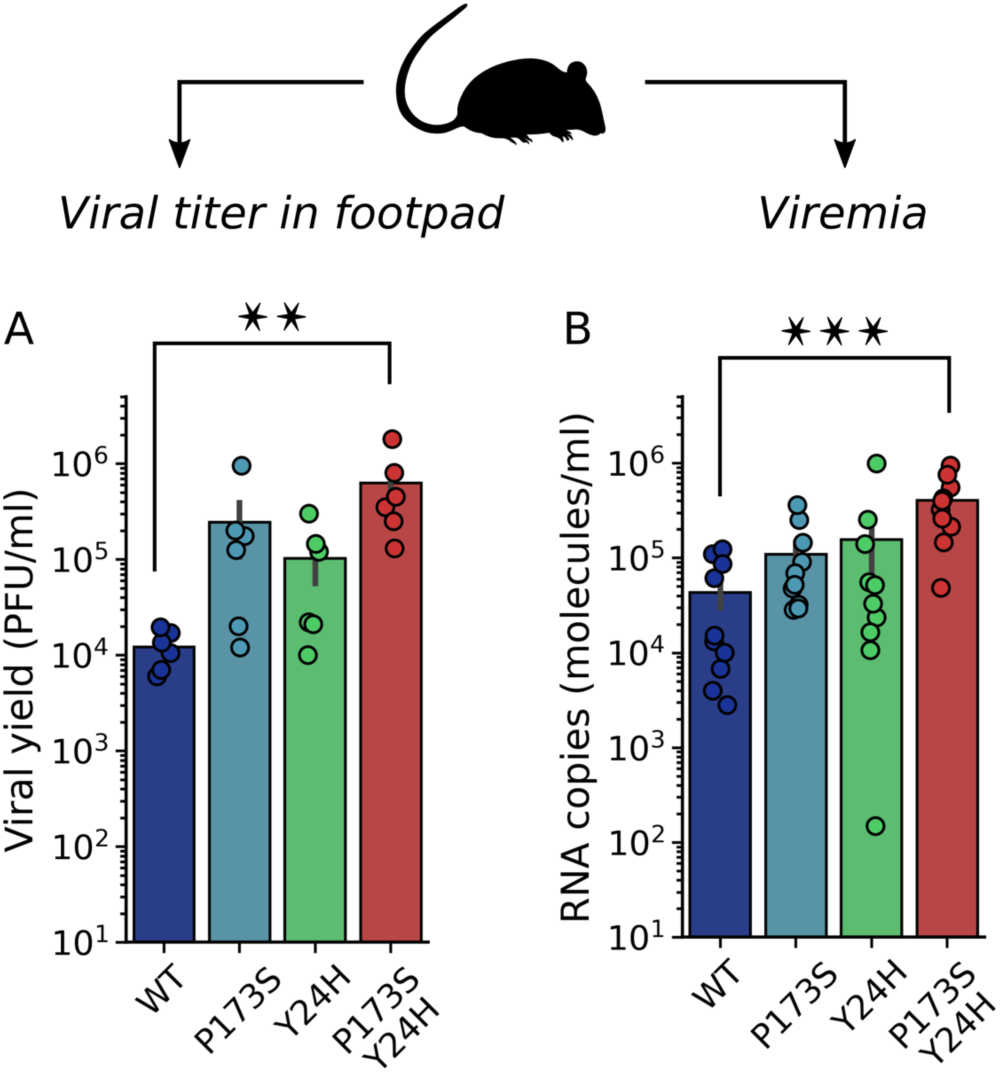
Infectivity of E1-Y24H and E2-P173S single and double mutants in mice. 4-5 week old male and female C57BL/6J mice were infected with 1000 PFU of WT or mutant CHIKV ZsGreen via subcutaneous inoculation in the footpad. Two days post infection: (A) the viral titer was determined in the footpad by plaque assay and (B) the viral load was quantified in the serum by qRT-PCR. For A and B, Kruskal-Wallis with Dunn’s post hoc test, * p < 0.05, ** p < 0.01, *** p < 0.001.

Next, we evaluated the fitness of mutant viruses in the vector host. First, we carried out growth curves in the mosquito C6/36 (*Ae. albopictus*) cell line (Figure 6A). Interestingly, we found that both E2-P173S and the double mutant reached a lower viral titer than WT (4-fold) and E1-Y24H showed a delay in the growth curve. In line with these results, mutant viruses also displayed decreased fitness in mosquitoes. Following feeding of *Ae. aegypti* mosquitoes with an infectious blood meal, all mutant viruses infected a lower percentage of mosquitoes than WT (Figure 6B), with a statistically significant reduction for E2-P173S and the double mutant. Interestingly, viral titers in the bodies of infected mosquitos were similar for the double mutant and WT (Figure 6C), suggesting a defect for the mutant virus to overcome the midgut infection barrier to establish an infection (22). In contrast, the viral titer for E2-P173S single mutant was two-fold lower than WT, suggesting an impact of the mutation in viral fitness in mosquitoes. Moreover, the result further shows an effect of E1-Y24H on the double mutant phenotype.

**Figure 6:**
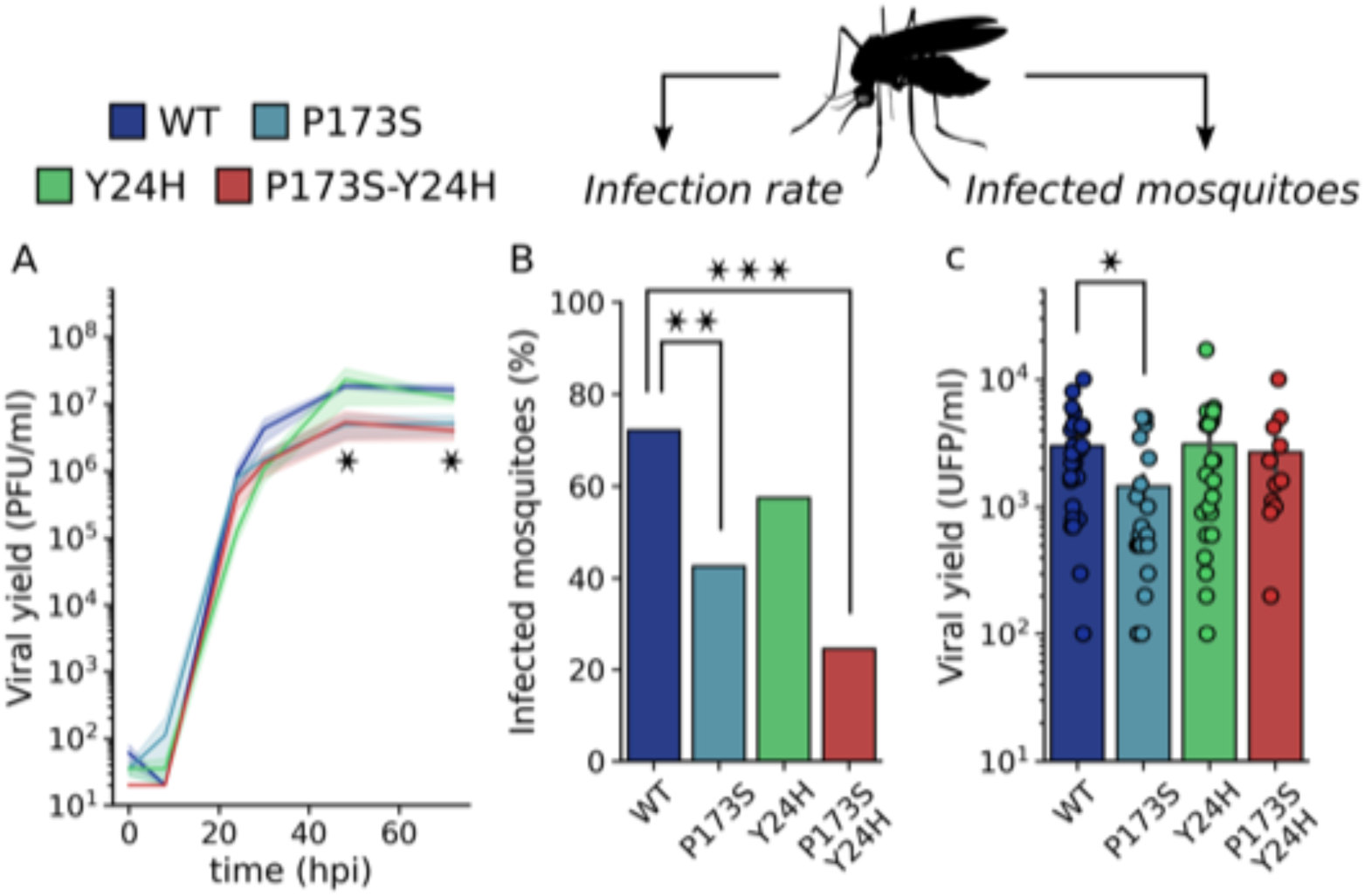
Infectivity of E1-Y24H and E2-P173S single and double mutants in mosquitoes. (A) Growth curve of WT and mutant viruses on C6/36 cells. Results represent the mean and standard error of three independent experiments. Kruskal-Wallis with Dunn’s post hoc test, * p < 0.05. (B and C) 4-7 day old female Ae. aegypti mosquitoes were infected with a bloodmeal containing 106 PFU/ml of WT or mutant CHIKV-ZsGreen. Seven days post infection the viral titer was determined in mosquito bodies. (B) Percentage of infected mosquitoes. Fisher exact test, * p < 0.05, ** p < 0.01, *** p < 0.001 (C) Viral yield in infected mosquitoes. Kruskal-Wallis with Dunn’s post hoc test, * p < 0.05.

Overall, these results show an opposite impact of E2-P173S and E1-Y24H on viral fitness in the alternate hosts, with a beneficial effect in mice and a detrimental effect in mosquitoes. In agreement with the *in vitro* characterization, the double mutant displayed a more pronounced phenotype than single mutants in both hosts. In addition, E1-Y24H partially rescued the deleterious effect of E2-P173S in mosquitoes, altogether reinforcing the functional interaction between these two residues.

### Impact of E2-P173S and E1-Y24H on viral particle stability and envelope proteins functionality

We hypothesized that mutations could impact protein structure and dynamics accounting for their effect on viral fitness. In this context, we next studied the thermal stability of the viral particle and the functionality of the envelope proteins of WT and mutant viruses. To study thermal stability, we quantified infectivity after incubation of viruses at 37°C in cell culture medium (Figure 7A). All viruses showed a similar stability profile, with approximately 25% of the initial infectivity retained after 8 hours of incubation and 2% after 24 hours indicating that mutations have no major impact on stability.

**Figure 7:**
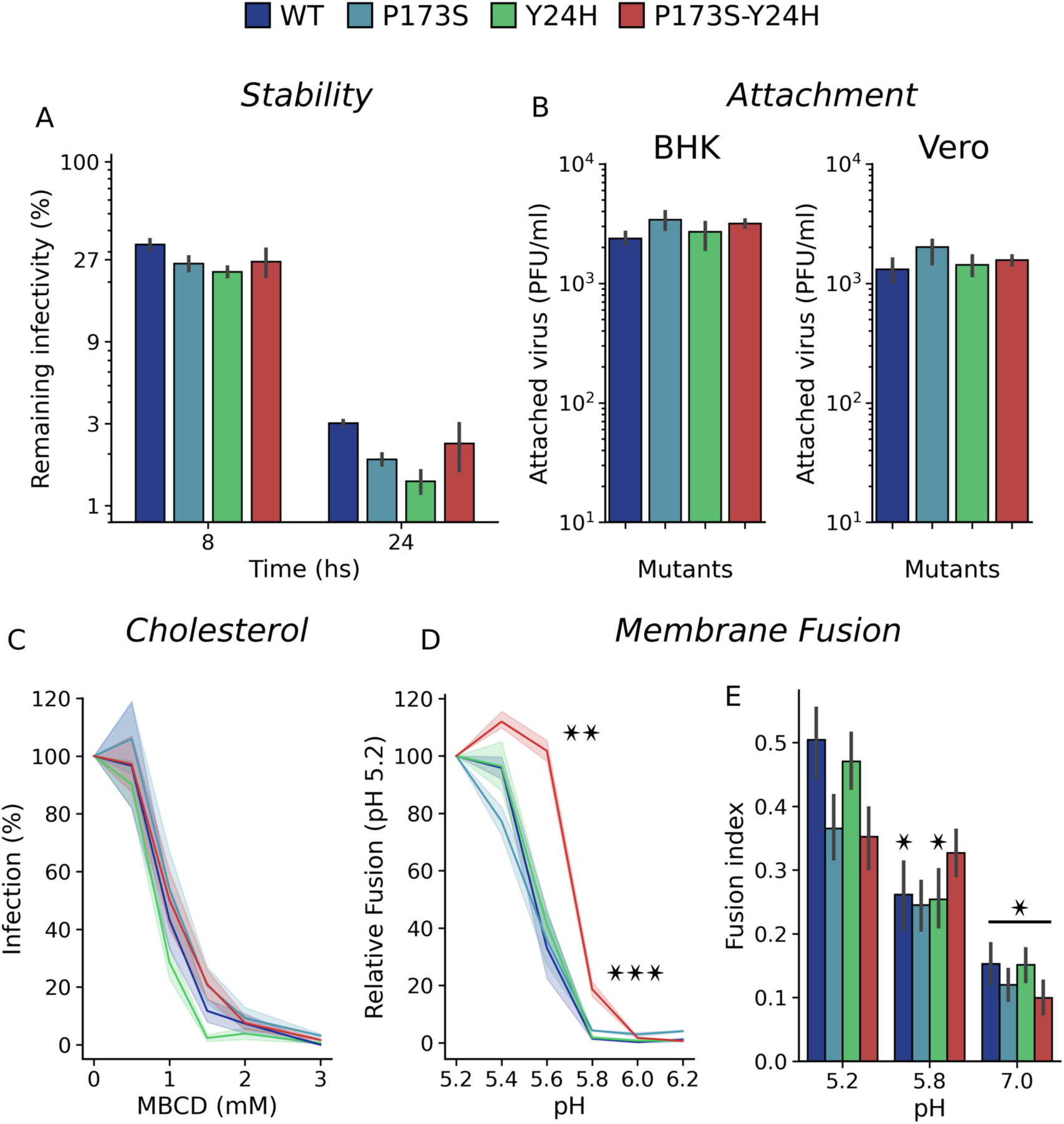
The impact of E1-Y24H and E2-P173S on envelope protein is associated with a shift in fusion pH threshold. (A) Thermostability of WT and mutant viruses. Viruses were incubated at 37°C in cell culture medium and the remaining infectivity after 8 or 24h was quantified using plaque assays. The figure displays virus titers at 8 or 24 hours compared to initial titers as percentages. (B) Attachment on BHK and Vero cells. BHK and Vero cells were incubated with WT and mutant viruses for 1 h at room temperature and the attached virus was quantified by plaque assay after cell lysis. Results represent the mean and standard error of the number of attached viruses for three independent experiments. (C) Cholesterol-dependence. BHK cells were treated with increasing concentrations of MβCD for 1 hour at 37°C and infected with WT or mutant CHIKV-ZsGreen viruses. Following infection, 20 mM NH4Cl was added to prevent further rounds of virus entry and the percentages of infected cells were measured one day post-infection by flow cytometry. Results represent the mean and standard error of the percentage of infected cells relative to the untreated control for three independent experiments. (D) FFWO assay. Membrane fusion was triggered by the treatment with PBS citric acid solution titrated at different pH. Results represent the mean and standard error of the percentage of infected cells relative to pH 5.2 for each virus for three independent experiments. One way ANOVA with Tukey’s post hoc test. Differences between each mutant and the WT for each pH, * p < 0.05, ** p < 0.01, *** p < 0.001. (E) Cell cell fusion assay. Results represent the mean and standard error of the Fusion Index of two independent experiments. One way ANOVA with Tukey’s post hoc test. Differences against pH 5.2 for each mutant, * p < 0.05.

Initial characterization of virus growth showing no differences in virus yields between WT and mutant viruses (Figure 3A) suggested that mutations do not alter CHIKV assembly or release. To further characterize the impact of mutations on protein functionality, we studied their effect on viral attachment, cholesterol-dependence for CHIKV entry, and fusion. Virus attachment assays showed no differences between mutant and WT viruses in either BHK or Vero cells (Figure 7B).

Target membrane cholesterol was shown to promote endocytic uptake of CHIKV and mutations in E1 were previously linked to an increased cholesterol dependency for fusion (6). To measure the cholesterol-dependance for viral infection, we used methyl-β-cyclodextrin (MβCD) to capture cholesterol and lower the levels in the plasma membrane prior to infection (23) (Figure 7C). All viruses were similarly sensitive to MβCD, indicating that the WT and mutant viruses display a similar cholesterol-dependance.

Finally, we assessed the fusogenic activity of WT and mutant envelope proteins. In the FFWO assay (Figure 7D left), the fusion degree against pH followed a sigmoidal curve with a marked fusion threshold for WT and mutant viruses. Both single mutants showed a similar fusion profile than WT virus. In contrast, there was a clear shift towards a higher pH in the threshold for fusion of E2-P173S E1-Y24H double mutant virus. We observed a similar behavior in the cell-cell fusion assay. While the fusion index at pH 5.2 was higher for the WT and E1-Y24H mutant, at pH 5.8 the fusion index was higher for the double mutant, indicating a shift of the fusion threshold towards neutral pH for this virus (Figure 7D right).

Altogether these results show that E1-Y24H and E2-P173S mutations qualitatively change the fusion phenotype, shifting the pH threshold for the double mutant, which confirms the functional interaction between these two residues.

### E2-P173S and E1-Y24H are located near two important hinges of CHIKV envelope glycoproteins

It is noteworthy that E2-P173S and E1-Y24H are apart from each other both in the E2-E1 heterodimer (99.2 Å) and in the viral trimeric spike (68.1 Å) (Figure 1). In order to gain deeper insights into the role of E2-P173 and E1-Y24 and the functional interaction between them, we performed molecular dynamics (MD) simulations of the envelope proteins in the pre-fusion conformation and constructed and analyzed the Residue Interaction Network (RIN) (29). To build the RIN, we computed the contact matrix and the Dynamic Cross Correlation Matrix (DCCM) of the Cα atoms from the MD trajectory (Figure 8A)(24). We used RIN to calculate residue communities using the Girvan-Newman algorithm (25) and mapped the resulting communities in the envelope proteins’ crystal structure. Communities represent groups of residues that move in a concerted manner. Detected communities were associated to the envelope protein domains but were not identical (Figure 8B and C).

**Figure 8:**
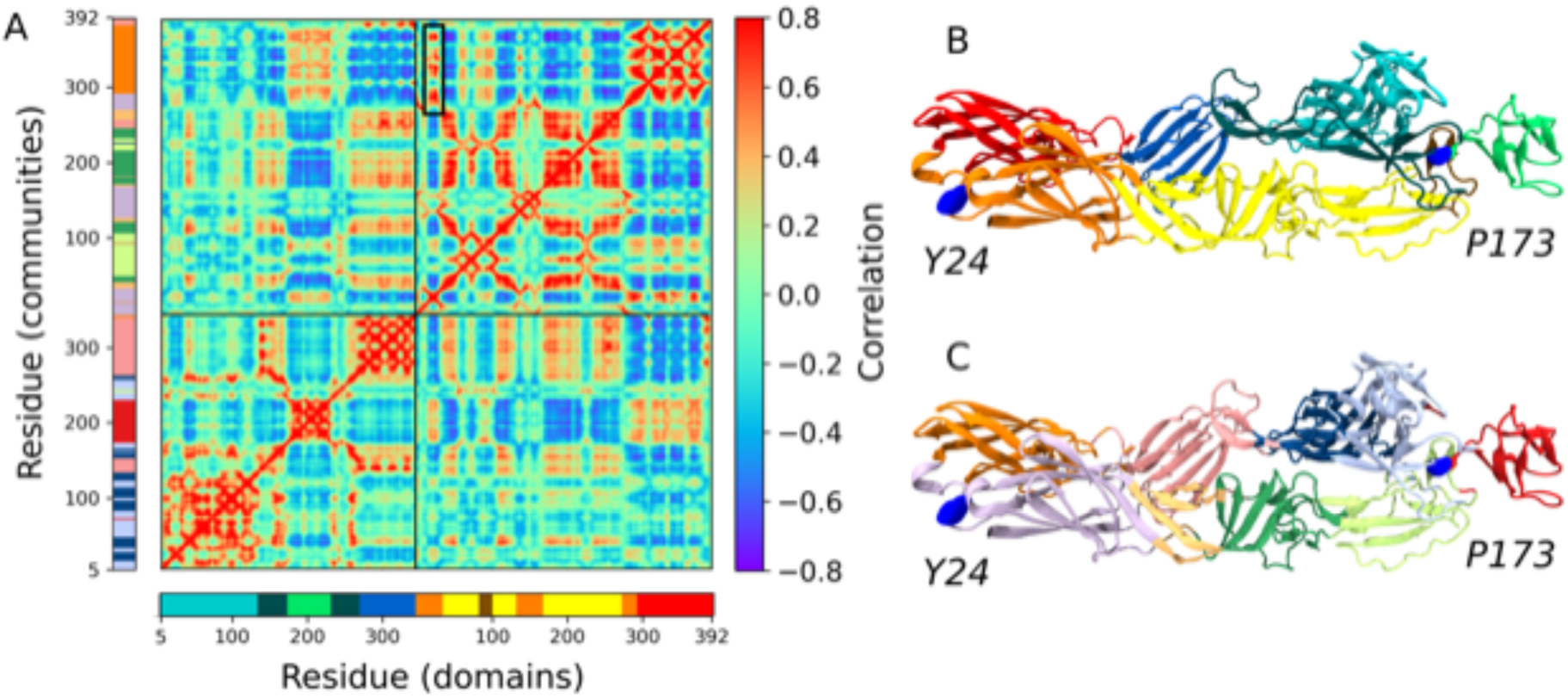
E2-E1 pre-fusion heterodimer structure and dynamics. (A) Dynamic Cross Correlation Matrix (DCCM) obtained from the molecular dynamics trajectory. The DCCM represents the correlation of motion between each pair of residues in the simulation. The black box shows the correlated motion between the E1-Y24 loop and domain E1-III. (B) Crystal structure of the envelope proteins of CHIKV (PDB ID 3N42) colored by domains: E1-I orange, E1-II yellow (fusion peptide ochre) E1-III red, E2-A cyan, E2-B green, E2-β-ribbon dark cyan and E2-C blue. E1-Y24 and E2-P173 are represented as blue spheres. (C) Envelope proteins colored by communities, detected by the Girvan-Newman algorithm form the Residue Interaction Network.

Next, we analyzed the position of each residue in the protein and in the RIN. E2-P173 is located between E2-B and the β-ribbon. This proline is adjacent to E2-P172, which is conserved in all alphaviruses (Supplemetary Figure 1). The PP motif is associated with a break in secondary structure. Additionally, in the context of the E1-E2 heterodimer these residues are located next to the arch 2 in the complementary strand of the β-ribbon (Figure 9A). The association between prolines and arches in the complementary strands of the β-ribbon is also present in arch 1 (E2-P269) and arch 3 (E2-P258 and E2-P260). Interestingly, these regions correspond to breakpoints between residue communities and, thus, represent breakpoints in residue connectivity (Figure 9A). In fact, E2-P173 is located at the hinge of the second Principal Component (PC) of the envelope proteins in a PCA of the MD trajectory, which is the PC associated with the highest RMSF of E2 domain B (Figure 9B and C). Overall, these data suggests that P172 and P173 would control the flexibility of the β-ribbon and E2 domain B, which is essential in the first steps of the conformational rearrangements of the envelope glycoproteins. Thus, E2-P173S may alter the behavior of the WT protein in this regard.

**Figure 9:**
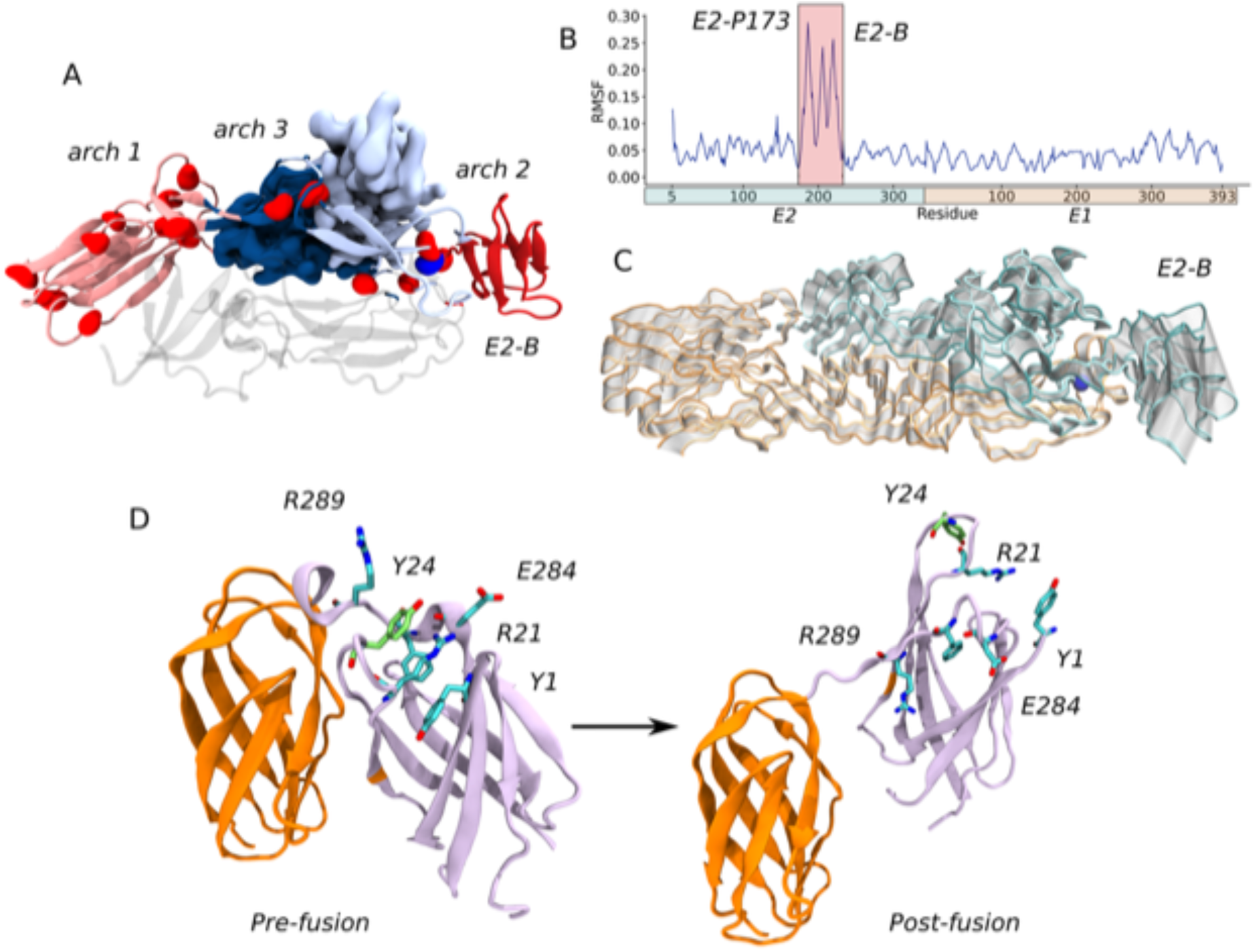
E2-P173 and E1-Y24 are located near flexible hinges in the envelope proteins of CHIKV. (A) E2-P173 is located in a hinge region in the β-ribbon. E2 domains C, B and the β- ribbon are represented as cartoons, E2-A is represented as surface and the tip of E1 is represented as transparent cartoons. Prolines on E2 are represented as red spheres (E2- P173 blue). The protein is colored according to MD communities. (B) RMSF per residue across the second Principal Component (PC) of a PCA of the MD trajectory. (C) Movement across the second PC. The protein is represented as tubes and the extreme conformations are colored in orange for E1 and cyan for E2. The conformations in between are represented as transparent tubes. (D) The E1-Y24 loop interacts with the flexible linker between domains E1- I and E1-III. Only domains E1-I and E1-III are represented for the pre-fusion (PDB 3N42) and post-fusion (PDB 1RER) conformations and colored according to MD communities. The residues involved in the interaction network with E1-Y24 in the pre-fusion conformation are represented as sticks, and are separated from each other in the post-fusion conformation.

In turn, in the MD simulations there was a strong interaction between the loop bearing E1-Y24 and the flexible linker between domains I and III (residues 283 to 294). Both loops were part of the same community (Figure 9D, pre-fusion) and showed a high correlation between each other and with domain E1-III (Figure 8A black box, table 1).

**Table 1:** E1-Y24 interaction network. Interaction energy and correlation of motion between Y24 and R21 with other E1 residues in the molecular dynamics simulations.

| Residue i | Residue j | $E_{\text{coul}}^{\text{A}}$ | $E_{\text{LJ}}^{\text{B}}$ | $E_{\text{total}}^{\text{C}}$ | Correlation<sup>D</sup> |
| --- | --- | --- | --- | --- | --- |
| Y24 | R21 | $-0.4 \pm 0.6$ | $-16.7 \pm 1.3$ | $-17.1 \pm 1.9$ | 0.906 |
| | P22 | $-2.2 \pm 0.2$ | $-6.1 \pm 0.2$ | $-8.3 \pm 0.4$ | 0.913 |
| | P26 | $-0.1 \pm 0.1$ | $-3.4 \pm 0.1$ | $-3.5 \pm 0.1$ | 0.914 |
| | E284 | $-10.8 \pm 3.4$ | $-5.5 \pm 0.6$ | $-16.2 \pm 4.0$ | 0.656 |
| | F287 | $0.5 \pm 0.2$ | $-10.2 \pm 0.2$ | $-9.7 \pm 0.4$ | 0.838 |
| | T288 | $-11.5 \pm 0.1$ | $-6.7 \pm 0.1$ | $-18.2 \pm 0.2$ | 0.873 |
| | R289 | $1.7 \pm 0.2$ | $-11.4 \pm 0.2$ | $-9.7 \pm 0.4$ | 0.938 |
| | V290 | $-1.0 \pm 0.1$ | $-6.9 \pm 0.1$ | $-7.9 \pm 0.1$ | 0.942 |
| R21 | Y1 | $0.3 \pm 0.2$ | $-14.0 \pm 1.6$ | $-13.7 \pm 1.8$ | 0.553 |
| | G23 | $-1.9 \pm 0.1$ | $-3.2 \pm 0.1$ | $-5.1 \pm 0.2$ | 0.919 |
| | Y24 | $-0.4 \pm 0.6$ | $-16.7 \pm 1.5$ | $-17.1 \pm 2.1$ | 0.906 |
| | S26 | $0.1 \pm 0.1$ | $-0.9 \pm 0.1$ | $-0.7 \pm 0.1$ | 0.842 |
| | E284 | $-70.7 \pm 6.8$ | $-0.2 \pm 1.0$ | $-70.9 \pm 7.8$ | 0.558 |
| | F287 | $-1.4 \pm 0.3$ | $-10.5 \pm 0.5$ | $-11.9 \pm 0.8$ | 0.69 |

(A) Coulomb interaction Energy. (B) Lenard-Jones interaction Energy. (C) Total interaction Energy. (D) Correlation of motion between residues.

E1-Y24 is an aromatic residue and establishes π-cation interactions with two nearby arginines (E1-R289 and E1-R21). In turn, E1-R21 interacts with E1-E284, E1-F287 and E1-Y1 building an interaction network that may be responsible for the correlated motion of the loop and the flexible linker. E1-Y24 loop is separated from E1-III and the flexible linker in the post-fusion conformation of E1 envelope protein (Figure 9D, post-fusion). This means that the interaction between the E1-Y24 loop and the linker has to break in the conformational change that drives the fusion process. Thus, the stability of this interaction may be important for the stability of the pre-fusion conformation of E1 and the regulation of the fusion process. At the acidic pH of the endosome histidine residues become protonated, thus the E1-Y24H substitution may impact on the strength of the interaction network connecting this residue to the linker in the context of the pH-triggered fusion process.

Altogether, MD simulations allowed us to propose a mechanistic hypothesis about the role of E1-Y24 and E2-P173 residues in the fusion process that could explain the molecular basis behind the interaction between E1-Y24H and E2-P173S mutations. While E2-P173 would be associated with the flexibility of domain B, E1-P173S may modulate the stability of E1 prefusion conformation. In this manner, both residues could be associated with important kinetic barriers in the conformational rearrangement of the envelope proteins and, thus, would play a role in the regulation of the timing and coordination of the different steps in the fusion process.

## Discussion

In this work, we studied the mechanism of action of a previously identified small molecule inhibitor of CHIKV infection and characterized the phenotype of viruses carrying mutations associated with antiviral resistance. Using complementary approaches, we demonstrated that compound 11 specifically inhibits the pH-dependent membrane fusion driven by CHIKV envelope proteins during entry. Characterization of mutant viruses indicated a functional interaction between E2-P173 and E1-Y24: viruses with the combined mutations E2-P173S and E1-Y24H displayed a distinct resistance to compound 11 likely associated to a shift in the fusion threshold towards higher pH, and showed a significantly increased replication in mice and impaired rate of infection in mosquitoes. We propose that E2-P173S and E1-Y24H cooperatively act to engage the heterodimer in its conformational rearrangement at a pH closer to neutral in comparison to WT.

### Functional interaction between E2-P173 and E1-Y24 in the regulation of the fusion process

The results presented in this manuscript support that E2-P173 and E1-Y24 work together in the regulation of the fusion process despite being distantly located in the envelope proteins of CHIKV. Interestingly, a connection between nearby residues has been previously observed. Infection of mosquitoes with a virus carrying a E1-V80Q mutation, close to E2-P173S, resulted in the selection of pseudorevertants with the second-site mutation E1-N20Y, near E1-Y24H (21). In the same line, the emergence of the epistatic mutations E1-K211T and E1-V156A in the context of the Indian Ocean lineage virus carrying a valine at E1-226 was associated with increased fusion at lower pH and higher sensitivity to NH_4_Cl treatment, suggesting a functional connection between residues 156, 211, and 226 in the entry dynamics (5). Taken together with our results indicating that compensatory mutations E2-P173S and E1-Y24H modulate the fusogenic function, these results suggest a link between two distant regions of the envelope complex.

Enveloped virus entry culminates with fusion of viral and cellular membranes and the delivery of the viral genome into the host cell cytosol. Virus envelope proteins anchored to the lipid envelope perform fusion following activation triggered by an external cue such as receptor binding or exposure to acidic pH. A common pathway for fusion involves protein conformational rearrangement from an initial prefusion state to the insertion of a fusion motif in the target membrane in an extended prehairpin state and final folding back into a postfusion hairpin state (26). Beyond the commonalities in the fusion process, virus fusion proteins diverge in structure and organization on the virus surface. Three canonical classes of fusion proteins are recognized based on their structure. Alphavirus E1 is a class II fusion protein. Cryo- electron tomography has recently allowed to visualize the pathway of CHIKV membrane fusion and together with cryo-electron microscopy and crystallography allowed to reconstruct conformational rearrangements of the mature E1-E2 heterodimer (9,26–28). Still details on the dynamics of molecular transitions at the residue level remain unresolved. Under acidic conditions, the central arch of E2 β- ribbon becomes disordered and domain B opens to expose the fusion loop at the tip of E1 (12,28,29). E2-P173 is located in a breakpoint in residue connectivity associated with the arch 2 of the β-ribbon and together with structural analyses our MD simulations indicate that this structure acts as hinge for the movement of domain B at the beginning of the fusion process. In turn, E2 dissociates from E1 that rearranges as a homotrimer in the post-fusion state. MD results showed that E1-Y24 loop strongly interacts with residues in the flexible linker between E1 domains I and III (E1-R289, E1-R21, E1-F287, E1-Y1 and E1-E284) in the pre-fusion conformation. Importantly, this interaction is broken during the conformational change of the envelope proteins towards the post-fusion state. Interestingly, E1-F287 and E1-R289 have been identified as a part of a different interaction network that stabilizes the post-fusion E1 trimer (30), suggesting that the conformational transition of the linker could be important in the regulation of the fusion process. Altogether, we hypothesize that E2- P173 and E1-Y24 are linked in the modulation of important kinetic barriers of the conformational change of the envelope proteins of CHIKV, acting as checkpoints of the fusion process. On the one hand, E2-P173S could alter the flexibility of the arch 2 of the β-ribbon, imposing a kinetic barrier to the opening of domain B. On the other hand, protonation of E1-H24 in E1-Y24H would lower the stability of the pre-fusion interaction network that stabilizes the linker, altering the energetic barrier that E1 needs to overcome in the transition between the pre-fusion and post-fusion states. The fact that the fusion phenotype is only altered in the double mutant, suggests a multistep reaction with sequential checkpoints. As the dissociation of E1 and E2, the assembly of the post-fusion trimer and the joint action of different trimers in the formation of the fusion pore require the coordination between different E1-E2 dimers in the viral particle (10,16), the relative rates of the different checkpoints may be fundamental for the coordination of the heterodimers.

### Impact of E2-P173S and E1-Y24H on *in vivo* infectivity

E2-P173S and E1-Y24H had opposite effects in fitness in experimental infection of mosquitoes and mice. They resulted in impaired infection rates in mosquitoes but enhanced viral loads in mice relative to WT virus. Further supporting a connection between these residues, in both mosquitoes and mice, the phenotype was more pronounced for the double mutant. In turn, both mutations arose after serial passages of CHIKV in presence of compound 11 in Vero cells (17), thus explaining the adaptive advantage in a mammalian host (31). Interestingly, the results highlight the evolutionary barrier imposed over viruses that alternate between hosts to develop resistance to an antiviral in the natural infection cycle (32). Mutations that confer resistance in the mammalian host to the antiviral are not necessarily adaptive in the alternate host and may even have a deleterious effect (33), preventing the fixation of the mutation in a natural population.

In further studies, it would be interesting to address the link between the change in the fusion phenotype, an increase in the fusion threshold, and the modulation of the viral fitness in the natural hosts. Notably, an opposite relationship between the fusion phenotype and the fitness in mosquitoes was observed in previous studies, where E1- V80L/Q mutants displayed a decreased fitness in mosquitos associated with a shift of the fusion threshold towards more acidic pH (20,21). These results suggest a complex relationship between the different functions of the fusion machinery that warrants further investigation.

In conclusion, in this study we characterized the mechanism of action of a small molecule inhibitor of CHIKV and showed that the compound inhibits the membrane fusion process during CHIKV entry. Additionally, through the study of resistance mutations to the antiviral activity of compound 11, we identified a functional interaction between two distant residues in the envelope proteins of the virus that, together, impact the fusion phenotype and the *in vivo* fitness of CHIKV.

## Materials and Methods

### Cell and viruses

Vero (Cercopithecus aethiops kidney, ATCC CCL-81) and HEK-293T cells (Human embryonic kidney cells expressing SV40 T antigen, ATCC CRL-3216) were grown in

Dulbecco’s modified Eagle’s medium (DMEM, Gibco) and BHK cells (Mesocricetus auratus hamster kidney, ATCC, CCL-10) were grown in MEM alpha medium (Gibco). All mammalian cell lines were grown at 37 °C in a 5% CO_2_ atmosphere in medium supplemented with 10% fetal bovine serum (FBS), 100 U/ml of penicillin and 100 μg/ml of streptomycin antibiotics (Gibco). C6/36 cells (*Aedes albopictus* larvae, ATCC CRL- 1660) were grown at 28 °C in Leibovitz L-15 medium (Gibco) supplemented with 10% FBS, 10% of Tryptose phosphate (29,5 g/l, Britania), 100 U/ml of penicillin, 100 μg/ml of streptomycin and 250 ng/ml of Amphotericin B antibiotics (Gibco).

WT and mutant CHIKV-ZsGreen of the Indian Ocean Lineage (Estearn Central and South Africa genotype) were derived from infectious cDNA clones as previously described . Viral titers of the final stocks were determined by plaque assay. Viral stocks were stored at −70 °C until use. The construction of E1-Y24H and E2-P173S single and double mutants is described elsewhere (17).

### Compound

The synthesis and purification of compound 11 was previously described (17). For biological assays, compound 11 was first dissolved in DMSO to a final concentration of 10 mM and then diluted in culture medium to the appropriate concentration. In all assays, the DMSO concentration was lower than 1%.

### Pseudovirus production and inhibition assay

The CHIKV ORF encoding for viral structural proteins was cloned into pCI-Neo (Promega) to obtain pCI-neo-CHIKV. HEK-293T cells at 70% confluence were co-transfected with plasmids psPAX2 (10 μg, Addgene #12260), pLB-GFP (10 μg, Addgene #11619), and pCI-neo-CHIKV or pMD2.G (2 μg, Addgene #12259) using polyethylenimine (PEI) reagent. Pseudoviruses were harvested two days post transfection and were concentrated by centrifugation at 3000 x g 4°C overnight.

For the determination of compound 11 inhibitory activity of pseudovirus infection, BHK cells at 50% confluence were treated with increasing concentrations of compound 11 and were infected with pseudoviruses (CHIKV or VSV) at a MOI of 0.05. 10 μg/ml of Polybrene (Sigma) was added to enhance pseudovirus adsorption. Three days after the infection, the percentage of infected cells was quantified by flow cytometry.

### Fusion from Without

Confluent BHK cells were treated with increasing concentrations of compound 11 and infected with CHIKV-ZsGreen WT or mutant viruses at a MOI of 0.1 at room temperature. Then, the inoculum was removed, and a PBS and citric acid solution titrated to an acidic pH with the corresponding concentration of compound 11 was added. After incubating for two minutes at room temperature, the acidic medium was removed, cells were washed with PBS, and α-MEM 20 mM NH_4_Cl was added. Cells were analyzed by flow cytometry at one day post-infection.

### Cell-cell fusion

BHK cells at 80% confluence grown on a coverslip were transfected with pCI-neo- CHIKV-GFP plasmid using PEI reagent. One day post transfection, cells were treated with increasing concentrations of compound 11 in culture medium for 40 minutes at room temperature. Subsequently, the medium was removed, and compound 11 was added in PBS titrated to an acidic pH with citric acid. After two minutes at room temperature, the acidic medium was removed, the cells were washed three times with PBS and incubated for 4 hours at 37°C before fixation with 4% PFA. Nuclei were stained with DAPI (4’,6-diamidino-2-phenylindole, Molecular Probes) and fluorescence microscopy images were captured using a Nikon Eclipse 80i Fluorescence microscope equipped with a DS-Qi1Mc camera with a 100X magnification.

Image analysis was followed to quantify syncytium formation in each treatment. Briefly, nuclei and GFP expressing cells were segmented in every image using local threshold and watershed algorithms. Afterwards, GFP expressing cells were manually classified in syncytium and not fused cells based on the number of nuclei and morphology. All cells with one nucleus were automatically considered an individual cell. For every image a fusion index was calculated as *1 - C/N*, where C is the number of GFP expressing cells and N the number of nuclei within GFP expressing cells in the image. Six random fields were taken per treatment and three independent experiments were performed.

### Plaque assay

Confluent Vero cells in 24-well plates were infected with serial dilutions of viral samples and were incubated for 1 h at 37 °C. Afterwards, 1 mL of overlay (DMEM 2% FBS, 0.4% methylcellulose) was added to each well. Three days post infection cells were fixed with 10% formaldehyde and were stained with crystal violet solution (20% ethanol, 0.1% crystal violet in water) to allow plate lysis count.

### Growth curves

Confluent Vero or C6/36 cells grown in 24 well plates were infected with WT or mutant CHIKV-ZsGreen at a MOI of 0.01. After incubating for 1 hour at 37°C, the inoculum was removed and cells were washed three times with PBS. 750 μl of culture medium was added, and the cells were incubated for three days at 37°C. At various time points, a 50 μl aliquot was removed, replaced with culture medium, and stored at -70°C. The viral titer at each time point was determined by plaque assay.

### Viral yield inhibition assay

Confluent Vero cells were treated with increasing concentrations of compound 11 and were infected with CHIKV-ZsGreen WT or mutant viruses at a MOI of 0.01. Cells were incubated for one hour at 37°C and then overlaid with culture medium containing the corresponding concentration of compound 11. Two days after the infection, the number of viral particles in the supernatant was titered by plaque assay.

### Competition assay

Virus stocks were diluted and mixed at 1:1 ratio to infect confluent Vero cells grown in 24-well plates at a MOI of 0.01. Supernatants of cells treated with compound 11 at 50 µM or mock-treated were collected three days after the infection and RNA was extracted using Quick RNA Viral kit (Zymo Research). The cDNA corresponding to the regions coding for E2 and E1 was obtained by RT-PCR using SuperScript III reverse transcriptase (Invitrogen) and primers 137 (5’-CGTTTGTAGATAACTGCGG-3’) and 95 (5’-TACTTAATTGTCGAGCTCTTAGTGCCTGCTGA-3’) for first strand synthesis of E2 and E1, respectively, and Pfx Accuprime polymerase (Invitrogen) and primers 136 (5’-GAAGAGTGGAGTCTTGCC-3’) and 137, and 93 (5’-GACTGAAGGGCTCGAGGTCA-3’) and 95 for PCR amplification of the corresponding fragments. PCR amplicons were sequenced by Sanger method and chromatograms analyzed using 4Peaks software (Nucleobytes).

### Experimental infection in mice

4-5 week old male and female C57BL/6J mice were infected subcutaneously in the footpad with 1000 PFU of WT or each of the mutant CHIKV-ZsGreen. At two days post infection, mice were euthanized to collect blood via cardiac puncture and to harvest the footpad. The footpad was ground in 1 mL of DMEM containing 2% FBS with steel beads using a Tissue-Lyser II (Qiagen) and debris was clarified by centrifugation at 8,000 x g for 10 minutes. Viral titers in the footpad were quantified by plaque assay on Vero cells. For serum, whole blood was centrifuged at 4,000 x g for 15 minutes and the serum was placed in Trizol. RNA was extracted following the manufacturer’s instruction and CHIKV genomes were quantified by RT-qPCR (Applied Biosystems RNA-to-Ct one step kit) with the following primers targeting CHIKV nsP4: 5’- TCACTCCCTGCTGGACTTGATAGA-3’ and 5’-TTGACGAACAGAGTTAGGAACATACC-3’, and probe: (5′-(6-carboxyfluorescein)- AGGTACGCGCTTCAAGTTCGGCG-(black-holequencher)]-3’). In vitro transcribed CHIKV RNA was used to generate a standard curve. All RT-qPCR samples were run in technical duplicates. All mouse experiments were performed in the Biosafety level 3 facility ABSL3 at the NYU Grossman School of Medicine in accordance with all Institutional Animal Care and Use Committee (IACUC) guidelines.

### Experimental infection in mosquitoes

4-7 days post-emergence, female Ae. aegypti mosquitoes (Poza Rica, Mexico; F40) were fed a blood meal containing 10^6^ PFU/mL of WT or each of the mutant CHIKV-ZsGreen supplemented with 5 mM ATP for 30 minutes. Engorged females were sorted and incubated at 28°C, 70% humidity, and 12h diurnal light cycle with 10% sucrose ad libitum for 7 days. After incubation, whole mosquitoes were ground in 300 ml of PBS with ceramic beads using a Tissue-Lyser II (Qiagen), and debris removed by centrifugation at 8,000 x g for 10 minutes. Viral titers in the bodies were quantified in the bodies by plaque assay. All mosquito studies were performed in the NYU Grossman School of Medicine ABSL3 facility.

### Viral particle stability

A 5x10^4^ PFU/ml dilution of WT or mutant CHIKV-ZsGreen was prepared in serum-free DMEM. At different time points (0, 8 and 24 hs) an aliquot was frozen at -70°C and the viral titer for each treatment was quantified by plaque assay.

### Viral attachment

Confluent BHK or Vero cells grown in 12 well plates were infected with WT or mutant CHIKV-ZsGreen at a MOI of 0.1. After incubating for 40 minutes at room temperature, cells were washed three times with PBS and were harvested in culture medium using a cell scraper. Cells were lysed by three cycles of freezing in liquid nitrogen and thawing at 37°C and the supernatant was clarified by centrifugation for 10 minutes at 1000 g and 4°C. The number of viral particles in the clarified supernatant was determined by plaque assay.

### Cholesterol dependence

A 25 mM MβCD (Thermo Fisher) solution in α-MEM with 25 mM HEPES at pH 7.5 was prepared for immediate use. Confluent BHK cells were treated with increasing concentrations of MβCD and were incubated for 1 hour at 37°C. Subsequently, cells were washed three times with PBS and were infected with CHIKV-ZsGreen WT or mutant viruses at a MOI of 0.1. After incubating for 1 hour at 37°C, the inoculum was removed, the cells were washed three times with PBS and medium containing 20 mM NH_4_Cl was added to prevent reinfection. One day post infection, the percentage of infected cells was determined by flow cytometry.

### Molecular dynamics and Residue Interaction Network

We performed molecular dynamics simulations of the CHIKV pre-fusion E1-E2 heterodimer in solution (PDB 3N42) using GROMACS 2021.2 (34). The protonation state at pH 7 of all protonable residues was defined using PROPKA (38). The Amber99SB*-ILDN force field (35,36) was used to describe the protein and a cutoff value of 10 Å was used for short-range electrostatic and van der Waals interactions. Long-range electrostatic interactions were treated with PME (Particle Mesh Ewald). The system was solvated using a dodecahedral box of TIP3P water (37) extending 12 Å from the protein’s surface. The system was neutralized with 0.15 M NaCl. After, the system was minimized, heated to 310 K and equilibrated for 200 ps in the NVT ensemble using the V-rescale thermostat (38) with a coupling constant of 0.1 ps. A second equilibration step of 1 ns was carried out in the NPT ensemble using the Berendsen barostat (39) with a reference pressure of 1 bar and a coupling constant of 2 ps. Finally, the production run was conducted in the NPT ensemble using the V-rescale thermostat and the Parrinello-Rahman barostat (40). In the equilibration and production runs, the LINCS algorithm (41) was used and a time step of 2 fs was employed. Four independent runs of 250 ns were carried out with the WT protein. For analysis, we concatenated the last 200 ns of each run. The molecular dynamics simulations were carried out on high-performance computing centers CCAD (https://ccad.unc.edu.ar/) and the High Performance Computing Portal at the NYU Grossman School of Medicine.

From the concatenated trajectory a PCA was conducted with GROMACS tools. The Residue Interaction Network (RIN) was built from the contact matrix and the Dynamic Cross-Correlation Matrix (DCCM) following the guidelines outlined in the work of Sethi and colleagues (24). The contact matrix was obtained using the CONAN tool (42). Two residues were considered to be in contact if they remained within 4.5 Å of each other for more than 75% of the dynamics. The DCCM was derived from the covariance matrix obtained in the PCA with GROMACS. The RIN was constructed using a Python script based on the NetworkX package (43). A node was defined for each residue, and an edge was established between two nodes if they were in contact during the dynamics. The weight of each edge was defined as the information transfer probability *dij* obtained from the correlation of each pair of residues in the dynamics (29). From the RIN, the betweenness centrality of each edge was calculated and the residue communities were identified using the Girvan-Newman algorithm (30) with NetworkX methods.

## Acknowledgements

We thank Felix Rey (Institut Pasteur, Paris) for critical reading of the manuscript. This work was supported by CONICET Doctoral Fellowships to L.B. and M.T.C. and funding from PEW Innovation Fund 2019 (D.E.A), Agencia de I+D+i PICT 2020-3371 (M.B. and D.E.A.), a start-up package from the NYU Grossman School of Medicine (K.A.S), and NIH/NIAID R01 AI162774-01A1 (K.A.S). D.E.A. and M.B. are members of CONICET.

---

**Supplementary Figure 1.**
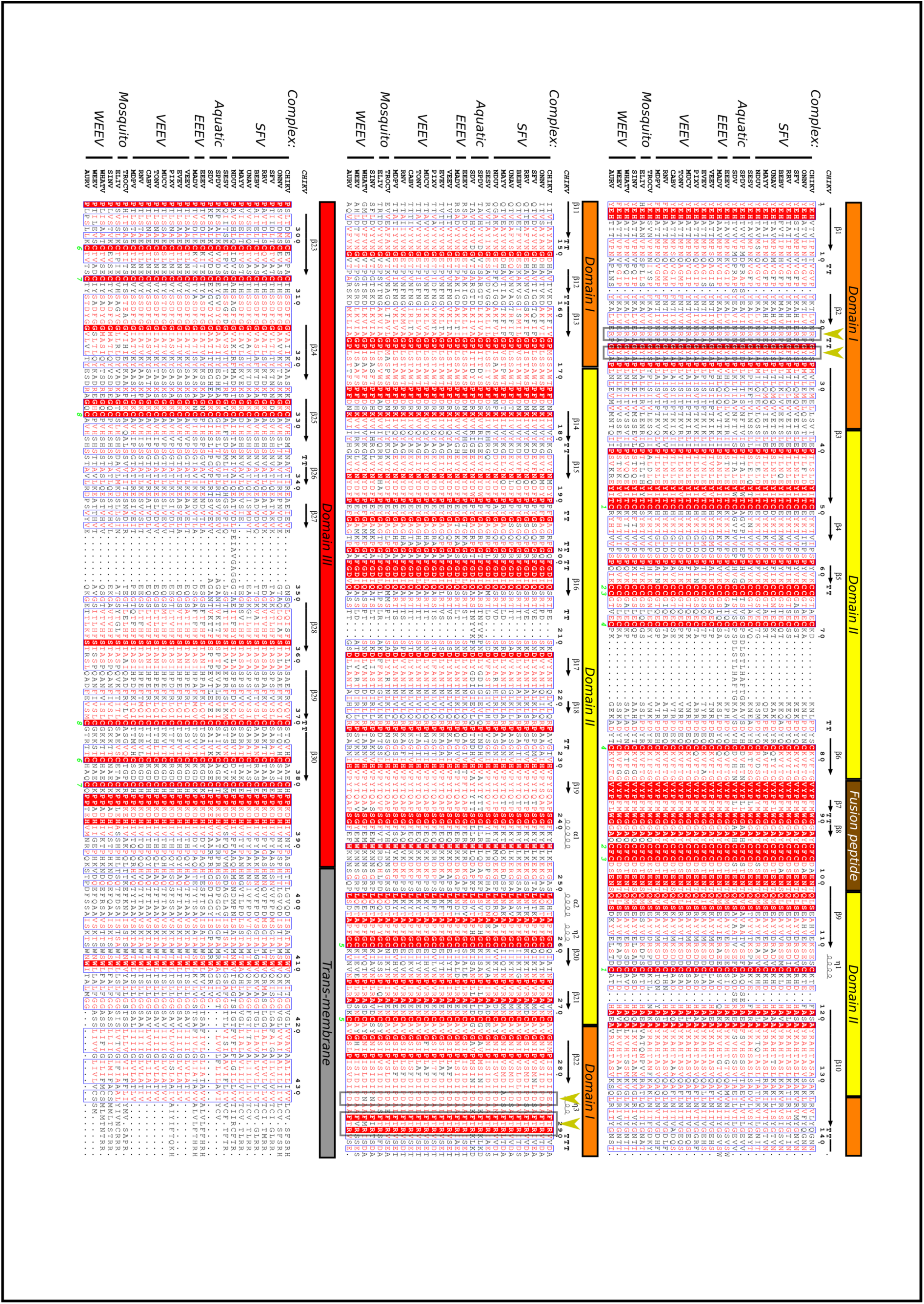
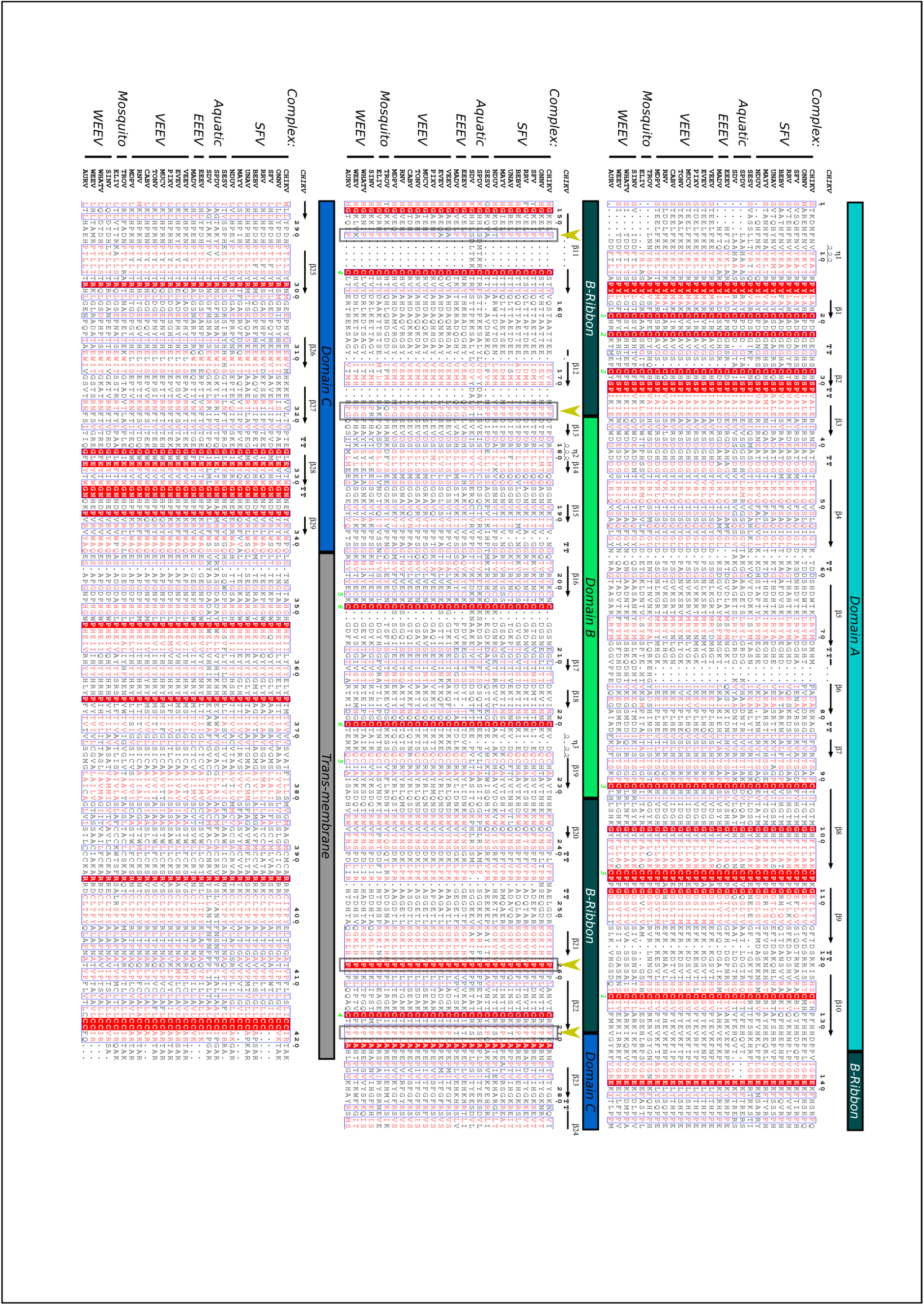
Amino acid alignment of alphavirus E1 and E2 envelope proteins. Alignments were generated using MAFFT (44) and visualized with ESPript (45). Strictly conserved residues are highlighted with a red background. Disulfide bonds are indicated with a green number below the corresponding cysteine residue. Conserved residues are displayed in red characters and boxed in blue. Protein domains depicted above the alignment are colored as defined in Figure 1 and symbols correspond to the secondary structure of the protein as determined in (9): arrows represent β-strands and coils represent either alpha helices (α) or 3/10 helices (η). Arrowheads in the alignment of E2 point to prolines associated with arches in the complementary strands of the β-ribbon. In the alignment of E1, arrowheads point to interacting residues between E1-Y24 loop and the flexible linker between domains E1-I and E1-III. Alphavirus complexes are indicated on the left as follows, Semliki Forest complex (SFV): Chikungunya virus IOL (CHIK IOL, JF274082.1), O’nyong-nyong virus (ONNV, NC_001512.1), Semliki forest virus (SFV, NC_003215.1), Ross River virus (RRV, GQ433354.1), Bebaru virus (BEBV, HM147985.1), Una virus (UNAV, HM147992.1), Mayaro virus (MAYV, NC_003417.1), Ndumu virus (NDUV, JX644166.1); aquatic virus complex (Aquatic): Southern elephant seal virus (SESV, NC_016960.1), Salmon pancreas disease virus (SPDV, NC_003930.1), Sleeping disease virus (SDV, NC_003433.1); Eastern Equine Encephalitis complex (EEEV): Eastern equine encephalitis virus (EEEV, NC_003899.1), Madariaga virus (MADV, KJ469622.1); Venezuelan Equine Encephalitis complex (VEEV): Venezuelan equine encephalitis virus (VEEV, NC_001449.1), Everglades virus (EVEV, NC_038671.1), Pixuna virus (PIXV, NC_038673.1), Mucambo virus (MUCV, AF075253), Tonate virus (TONV, NC_038675.1), Cabassou virus (CABV, NC_038670.1), Rio Negro virus (RNV, NC_038674.1), Mosso das Pedras virus (MDPV, NC_038857.1); mosquito-specific virus complex (Mosquito): Trocara virus (TROV, NC_043402.1), Eilat virus (EILV, NC_018615.1); Western Equine Encephalitis complex (WEEV): Sindbis virus (SINV, NC_001547.1), Whataroa virus (WHATV, NC_016961.1), Western equine encephalomyelitis virus (WEEV, NC_003908.1), Aura virus (AURV, NC_003900.1).

